# MEMS enabled miniaturized light-sheet microscopy with all optical control

**DOI:** 10.1101/2021.02.13.431066

**Authors:** Spyridon Bakas, Deepak Uttamchandani, Hiroshi Toshiyoshi, Ralf Bauer

## Abstract

We have designed and implemented a compact, cost-efficient miniaturised light-sheet microscopy system based on optical Microelectromechanical Systems (MEMS) scanners and tunable lenses. The system occupies a footprint of 20 × 28 × 13 cm^3^ and combines off-the-shelf optics and optomechanics with 3D-printed structural and optical elements, and an economically costed objective lens, excitation laser and camera. All-optical volume scanning enables imaging of 340 × 190 × 60 µm^3^ volumes with 0.25 vps and minimum lateral and axial resolution of 0.9 µm and 2.95 µm respectively. An open-top geometry allows imaging of samples on flat bottomed holders, allowing integration with microfluidic devices, multi-well plates and slide mounted samples, with applications envisaged in biomedical research and pre-clinical settings.

## Introduction

Over the last decade, light-sheet microscopy (LSM) has established itself in the fluorescence microscopy field as a fast 3D imaging technique with reduced background signal and lower photobleaching rates compared to the most commonly used approaches such as widefield fluorescence microscopy and confocal microscopy. These features originate from a spatial de-coupling of fluorescence excitation and imaging by using a sheet excitation of fluorescence in the focal plane of the imaging objective, allowing a full field read-out while exciting only a thin slice of a sample^1–3^. In general, LSM variants employ an excitation sheet generation through either cylindrical lenses or linear scanning of Gaussian or non-diffracting beams. Cylindrical lenses focus a Gaussian beam in only one axis, creating a reshaped thin slice of light in a design known as selective plane illumination microscopy (SPIM)^1,3,4^. Even though SPIM and its derivatives have shown good results for various imaging questions^4,5^, they suffer from non-uniformity of the intensity profile of the light-sheet due to the Gaussian beam profile and allow less control over the excitation beam. Creating the light-sheet through scanned excitation is known as digital scanned light-sheet microscopy (DSLM) in its original representation and introduces a pivoting mirror in the illumination path^2,3,6^. The excitation beam gets reflected off a fast-rotating mirror, resulting in a scan line which is then focused down to the sample area. With this method the light-sheet can achieve a much more uniform intensity profile that can be tailored to the desired field of view (FOV) with the control of the mirror. These initial demonstrations have opened the door for a vast, growing field of system demonstrations for increasing imaging speed, resolution and adaptability, with notable examples being work on using non-diffracting beams for thinner light-sheets with wider FOV^7,8^, super-resolution approaches to improve specifically axial resolution limits from light-sheets^9,10^, single objective light-sheet approaches such as Oblique Plane Microscopy^11^, SCAPE^12^ or SOPi^13^ and multi-photon approaches^14^. In most variations, z-stacks are acquired by stage scanning approaches to create 3D data volumes, allowing for a pre-determined fixed alignment position between the two orthogonal excitation and detection light paths. This however limits imaging speeds to avoid movement artefacts, specifically in non-fixed samples. This limitation has been circumvented to an extent with mirror based volume scanning in SCAPE and adaptive optics elements for accessing different 3D image planes^15,16^. A further limitation requiring, in most LSM systems, special sample holders due to the spatial constraints of orthogonal objectives has in recent years been addressed by moving to open-top configurations with solid or water based immersion lens geometries^17–19^ or the mentioned single objective variants ^11–13^ to allow investigations using samples in standard microscope slides and cover slips, cell dishes or well-plates.

Even though LSM has established itself as one of the main tools in fluorescence imaging for larger biomedical samples, its use can be limited due to the high cost for designing or purchasing an LSM imaging system. Additional limitations occur due to most LSM approaches targeting specific applications. As a result, the designs are not easily transferable between different studies, where a complete reconsideration of sample holders and optical path adjustment needs to be made. A few attempts have managed to make LSM more accessible by providing self-assembling protocols such as open-SPIM^20,21^ or plug-ins for conventional microscopes^22^. Specifically, the open access microscope design guide and detailed listing of the optical elements needed in the practical implementation of the imaging system is allowing access to a wide range of potential users, with the cost difference compared to the commercial solutions of DLSM and SPIM within one order of magnitude.

Building on the range of LSM variants described above, and recognising the need to make systems even more widely available in lower cost settings and settings with space constraint, we present a compact and low-cost version of an LSM microscope adopting the advantages of a flexible DSLM methodology with the use of Microelectromechanical Systems (MEMS) technology as active alignment and control elements^23,24^. The LSM implementation presented has all optical positioning of both orthogonal beam paths, without the need for stage scanning for 3D volume data generation of up to 0.25vps. A two-axis MEMS micromirror^25^ is used for control of the light-sheet excitation beam, allowing a 37 kHz fast resonance mode to create the scanned light-sheet and an orthogonal static displacement mode to address the parallel planes needed for 3D reconstruction of the sample. The use of a small scale electrically tunable lens in the imaging path allows for a synchronised focal plane adaptation of the full-field fluorescence collection. In this way we introduce a miniaturized mechanical-movement-free LSM, reducing the cost and bulk of the system to £5k and 20×28×13 cm^3^, as well as providing full digital control of all optical paths. Standard sample mounting techniques on cover slips or flat-bottomed well-plates can be used due to the integration of custom 3D-printed optical prisms for beam coupling and optical aberration reduction. The optical design and choice of elements in our miniaturized LSM functions as a cost and size effective solution for versatile and compact use for a vast variety of imaging questions ranging from cancer and neuroscience applications in microfluidic devices over investigations in antimicrobial resistance build-up to potential histopathology applications.

## Results

### MEMS light-sheet microscope configuration

The presented miniaturized LSM system design (Figure 1 (a) and 1 (b)) is based on a DSLM methodology where the physical footprint reduction and low-cost construction aim is achieved through the active MEMS elements with a balanced trade-off between imaging FOV and lateral and axial resolution. Off-the-shelf small diameter achromatic lenses are used for shaping the scanned light-sheet excitation, with a cost efficient x20, 0.4 NA objective providing a balance between imaging performance, size and working distance. Sample mounting using standard cover-slips or well plates is achieved through air coupled objective lenses and a 3D-printed solid immersion prism. The prism has a 30° angle between the imaging normal and sample normal in order to allow physical positioning of large samples and of the optical elements.

**Figure 1.**
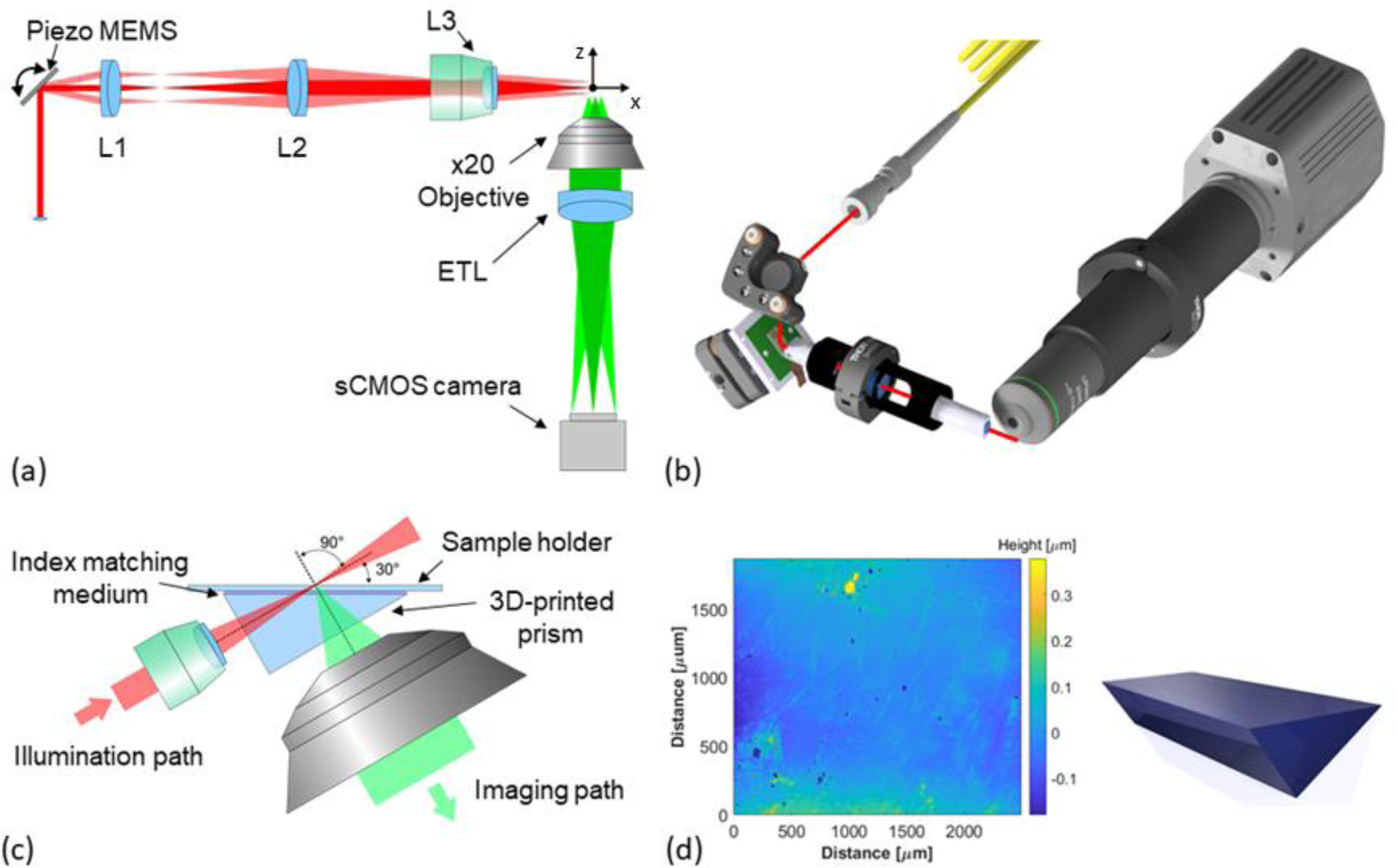
Microscope Optical setup. (a) Miniaturized LSM top view with illumination path (red) and imaging path (green). The collimated excitation beam is resonantly scanned out of plane (y-axis) to create the light-sheet and positioned in-plane (z-axis) to create all optical 3D image scanning using a MEMS micromirror. The achromat excitation lenses (L1-L3) create a telecentric system in sample space. An electrical tunable lens (ETL) allows fast focal plane synchronisation with the excitation. (b) 3D representation of the miniaturized LSM, using a mix of custom 3D-printed mounts and of-the-shelf lens tubes for a robust, compact and inexpensive device approach. The excitation beam is guided onto the scanner in the plane of the scanned light-sheet to avoid scanning induced field curvature. (c) Schematic of the optical beam paths at the sample, including an optical clear 3D-printed prism to reduce aberrations for imaging of specimens on flat sample holders. (d) Measurement of the surface profile of the 3D-printed prism with overall roughness in the sub 100nm range.

The light-sheet is generated using a 473 nm laser diode, which is fibre coupled for single mode operation and flexible deployment at multiple miniaturized LSM setups. A collimated Gaussian beam with full-width half-maximum (FWHM) of 400 μm is guided to a 45° positioned 2D piezoelectric (PZ) scanning MEMS mirror which creates the light-sheet and orthogonally positions it in the 3D sample space. The scanned beam gets 4x magnified through a 4f lens system and focused into the sample with a single achromatic lens with effective NA of 0.15. The beam path with the MEMS mirror at rest results in the beam being focused at an x-distance of 5.2 mm from the focusing lens. Resonant rotation of the mirror generates a height controllable focused scan line with FWHM beam waist of 2.95 µm. The optics are housed in a half inch diameter aluminium tube, with custom 3D-printed adapter holders for accurate assembly and positioning. The scanning MEMS mirror is housed in a 3D-printed holder specially designed to sit at a 45° angle between the two parts of the illumination path.

The imaging path is designed around the mentioned x20 0.4NA objective with finite correction at 160 mm mounted in a one-inch lens tube system. The objective allows for positioning at a working distance of 8.72 mm orthogonal to the focusing plane of the scanned illumination beam. An electrically tunable lens allows control of the focal position in the sample without the need for stage movement. Within the addressable positioning range of the imaging volume, the lens does not introduce significant aberrations over the field of view. It is placed directly after the objective with the use of 3D-printed adapters. The aperture of the electrically tunable lens generates an effective NA of the imaging arm of 0.28. A long pass filter with 500 nm cut off wavelength is placed before the sCMOS camera in order to block scattered excitation light. The total measured footprint of the microscope is 16 cm by 20 cm in its current configuration.

The microscope geometry requires microscope slides to be positioned at a 30° angle towards the imaging path to enable imaging due to the geometrical constrains of the LSM configuration. For this reason, we introduce a custom designed 3D-printed prism placed in the area between the imaging objective and the illumination lens closely coupled to the cover slip (Figure 1 (c)). The prism creates flat interfaces for both beam paths with matched 30° angle towards the sample and is coupled to glass bottomed slides or dishes using index matching oil in order to reduce aberrations. The prism is 3D-printed on a consumer printer using a clear resin with a refractive index of 1.53. A spin coating post-processing step of the optical surfaces of the prism results in surface roughness of <100 nm (see Figure 1 (d)), with high optical transmission.

The working principle of the miniaturized LSM microscope is based on the capabilities of the 2D scanning MEMS. More specifically, the scan line is generated with a sinusoidal waveform input at the resonance frequency of the mirror, which results in a telecentric light-sheet with thickness equal to the 2.95 µm FHWM of the Gaussian excitation. The static rotation of the scanning MEMS can translate the light-sheet in parallel planes along the z-axis relative to the rest position. This movement is synchronised with tunable lens changes in the imaging path to create the 3D sample content, with all active elements controlled via LabView and an Arduino microcontroller for synchronisation and timing. The synchronization approach used for the two controllable MEMS elements follows a combinational linear step changes of the two elements starting from the maximum +z and incrementing towards the final -z position. The collected stack of images represents the whole 3D voxel with dimensions of 340 μm height (y), 190 μm length (x) and 60 μm depth (z), defined by the imaging path FOV and maximum light-sheet translation distance.

### PZ Scanning MEMS and tunable lens

The piezoelectrically actuated silicon-on-insulator MEMS (Figure 2 (a))^25^ with 1.1 mm mirror diameter provides full control of the excitation beam position in two axes. The thin-film piezoelectric PZT material layer deposited on the patterned silicon chip is responsible for enabling mechanical actuation, with in-plane stresses being transferred to angular mirror rotations through the device geometry. The device features two different actuation methodologies for the two movement axes, enabled by two oppositely placed serpentine springs for the static rotation and an inner ring frame for the fast resonance pivoting. Specifically, for the static rotation of the mirror the actuators on the serpentine springs are divided into pair-wise actuators with one part each responsible for an opposite static tilt movement per section. The sequential actuation of each section group creates the overall positive and negative tilt of the mirror frame and with it the mirror around the x-axis. The fast pivoting rotation is enabled with a sinusoidal waveform input to the inner ring frame actuators, which allows resonance excitation of the mirror rotation eigenmodes. This is based on the geometry of straight rectangular suspension beams between the mirror and the frame, which allows resonant rotation around the y-axis.

**Figure 2.**
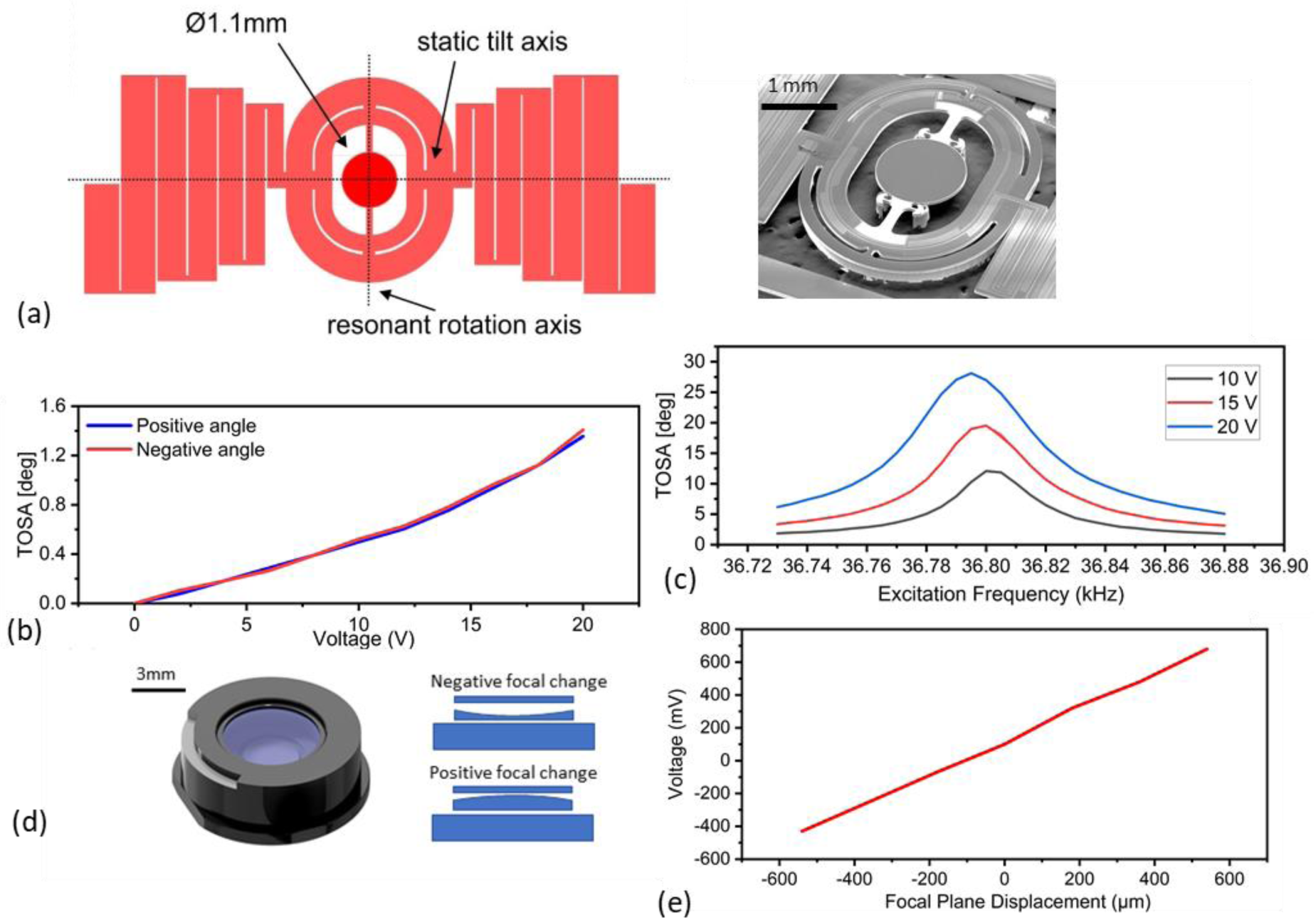
Characterisation of the optical MEMS devices. (a) Schematic and SEM Image of the 2D PZ scanning mirror featuring spiral actuators for static rotation and an inner ring frame for fast resonance rotation. (b) Total optical scan angle (TOSA) characteristics near the fast axis (scan along y-axis) resonance peak of 36.8 kHz for three different voltage inputs of 10 V, 15 V and 20 V (max). (c) Total optical scan angles of static rotation (scan along z-axis) with voltage input up to 20 V (max) for positive and negative angles. (d) Schematic and working principle of commercial MEMS tunable lens Optotune EL-3-10. (e) Tunable lens focal shift characteristic in the miniaturized LSM system.

The PZ MEMS mirror characteristics show a resonance tilt movement of the mirror around 36.8 kHz, with the full optical scan angle being measured for three different actuation voltages and reaching a maximum peak of 27° at 20 V_pp_ (Figure 2 (c)). Furthermore, the static angle along the orthogonal axis allows positioning of the mirror with a maximum of 1.5° optical scan angle for each movement direction at a maximum voltage of 20V (Figure 2 (b)). The response time needed between small signal voltage steps during static actuation is measured as <1ms to estimate image acquisition dead-times if a step wise image acquisition scheme is used.

The active element responsible for adjusting the imaging focal plane position is an Optotune EL-3-10 electrically tunable lens (Figure 2 (d)) through its shape changing technology. The lens can address positive and negative focal length changes with a clear aperture of 3 mm. The resulting focal plane shift in the presented LSM system is illustrated in Figure 2 (e) with the total addressable axial position change exceeding 400 μm as the lens is driven with operating voltages from −0.2 V to 0.2 V. Small step rise times for curvature changes are <1 ms, with settling times in the low single digit millisecond range as well.

### Illumination path characterisation

Characterization of the LSM illumination path is accomplished using the fluorescence response in the xy-plane through a fluorescein solution filled cuvette, by imaging the non-scanned beam propagation through a 0.4 NA objective along the x-axis for varying z positions (Figure 3 (a)). Images are acquired for the centre position and maximum and minimum static angles of the PZ MEMS, resulting in three different propagation positions of the illuminating beam. The beam diameter is measured throughout the propagation through the FOV, with selected cross-sections showing the Gaussian beam profile at the beam waist and edges of the LSM imaging FOV (Figure 3 (b)). A FWHM waist equal to 2.95 μm is measured with a confocal parameter of 74 μm (Figure 3 (c)), which leads to a beam diameter increase to 8.5 µm at the edge of the imaging FOV. The waist diameter stays constant for the different z-axis positions, with a maximum shift of the waist position along the propagation axis of 1 µm.

**Figure 3.**
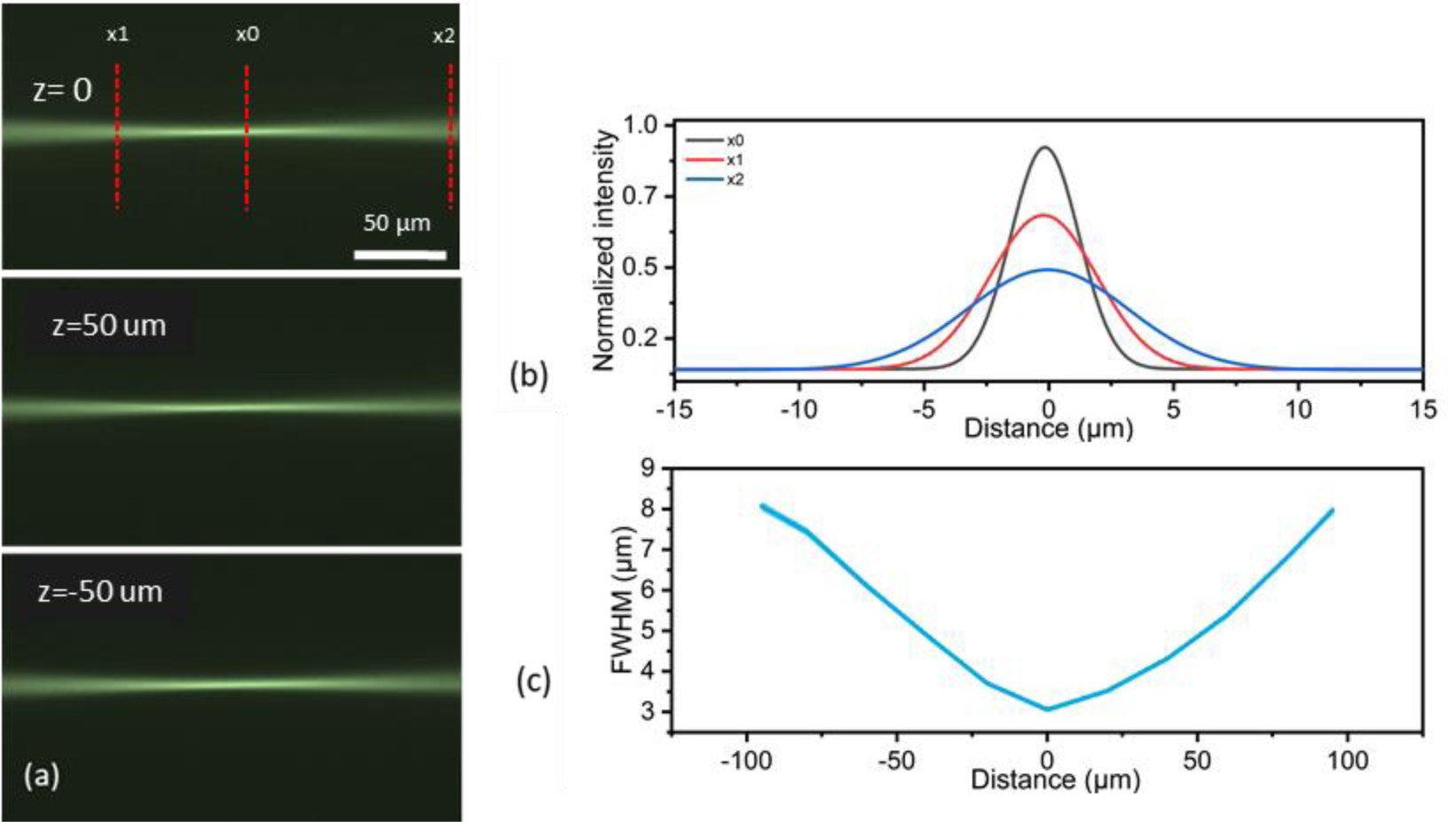
Excitation beam path characterisation. (a) Propagation through a fluorescein cuvette at z=0 (unactuated), z= 50 μm (maximum negative actuation) and x = −50 μm (maximum positive actuation). (b) Normalized intensity graphs are presented along the length of the beam for three different positions x1= −50 μm, x0 = 0 μm, x2 = 95 μm. (c) Beam width propagation through the x-axis with beam waist of 2.95 μm FWHM.

**Figure 4.**
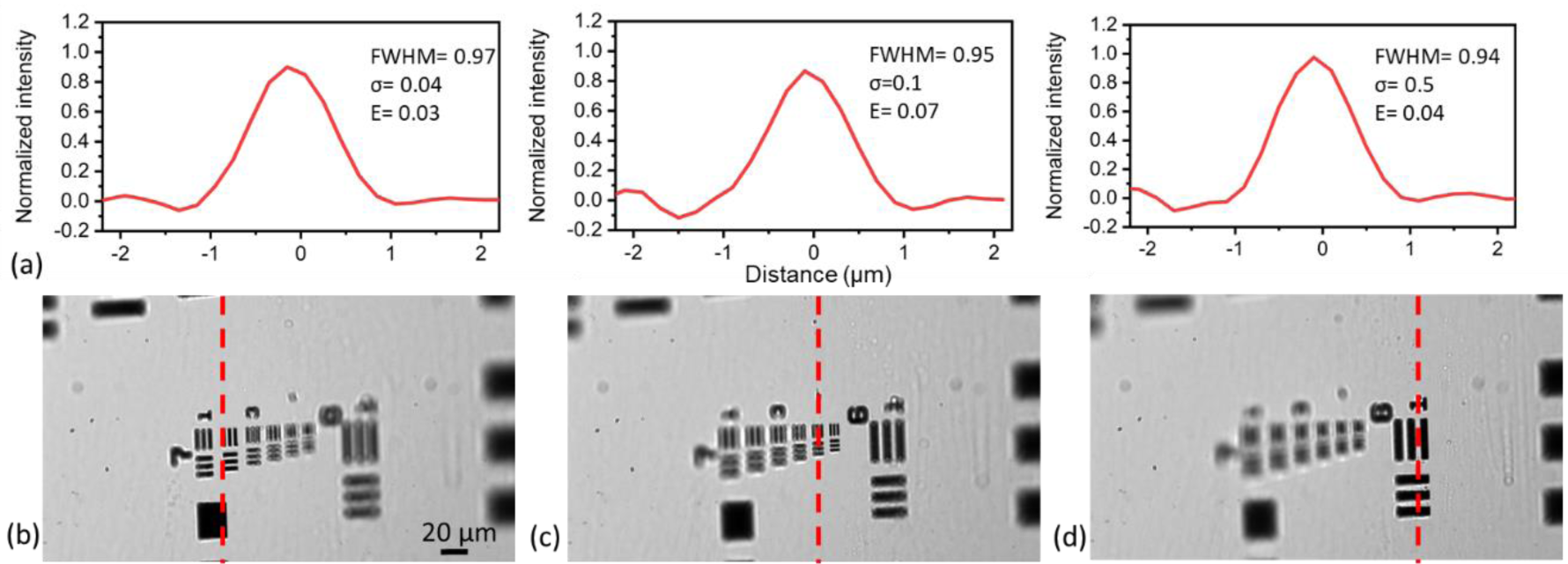
Imaging path characterisation. (a) Experimental PSFs derived from the Edge Spread Function for all three focal plane positions, showing a resolution limit of 0.95μm FWHM. (b) Cropped image of 30° USAF 1951 target group 7 for three different inputs of the tunable lens. Dotted vertical lines indicate where the focal point of the image is centred. z= −50 μm with Vtunable = −60 mV, (c) z = 0 μm with Vtunable = 0 V, (d) z = 50 μm with Vtunable = 60mV.

The characterization of the light-sheet is achieved with actuating the fast-rotating mode of the PZ MEMS at the resonance frequency of 36.8 kHz and imaging this time the propagation beam along the y-axis. The light-sheet can reach a maximum height of 550 μm with a voltage input of 20 V and sheet waist width equal to the beam waist FWHM of 2.95 μm.

### Imaging path characterisation

The imaging path analysis uses a USAF 1951 chrome plated target mounted at a 30° orientation to the imaging path normal, identical to the angle used for microscope slide mounted samples. Following the slanted edge method, the Line Spread Function (LSF) (Figure 4 (a)) is experimentally measured for the centre (Figure 4 (c)) and edges (Figure 4(b) and 4(d)) of the FOV, based on measurements of group 7 elements of the imaging target with a resulting resolution of 0.95 μm throughout the imaging depth defined by the angled sample. The FOV using the 2/3” sCMOS camera is measured at 190 μm by 340 μm with the tunable lens at its unactuated position. Additionally, the change in focal plane is characterized for the full range of the tunable lens actuation, with imaging depths changes of 60 μm achieved using a 70 mV actuation. This depth is matching the total light-sheet orthogonal positioning travel set by the PZ scanning mirror. The use of less than 4% of the addressable tuning range of the lens limits spherical aberrations introduced by the lens change.

**Figure 5:**
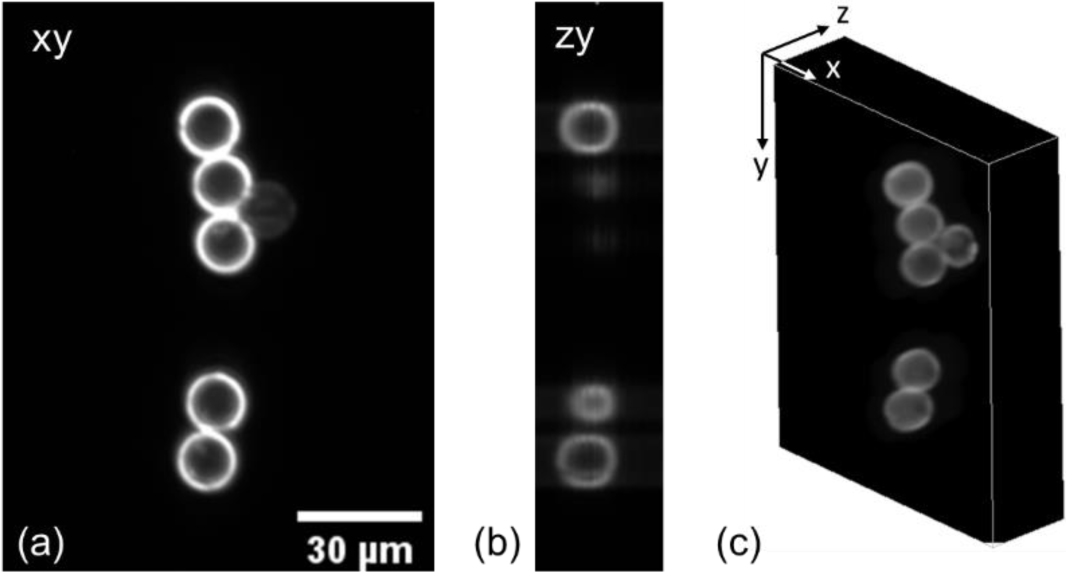
Sections and volume view of 15 µm surface stained microbeads. (a) xy slice of image volume, showing clear rings with negligible background, (b) zy section (c) volume view of surfaces

### Imaging of fixed nanobeads

Fluorescence labelled sub-resolution nanobeads and microbeads are imaged using 473 nm excitation. While beads of <1 µm size are fully labelled throughout the volume and used for point spread function evaluation, beads with 6 µm and 15 µm size are ring stained and used for evaluating sectioning ability of the light-sheet. A fixed microbead slide was used for the sectioning evaluation, with an exemplar image slice shown in Figure 5 (a) and 5 (b) and a 3D representation in Figure 5 (c). The MEMS and tunable lens were synchronised and moved in Nyquist limited 0.6 µm steps through the sample volume, with a laser power of 0.3 mW, camera exposure time of 70 ms, 2 by 2 binning and 50 steps to image a volume of 100 × 135 × 30 µm^3^. Volume acquisition is completed with 0.25 vps. The resulting images for a cropped slice show clear sectioning ability with low to non-existent background.

**Figure 6.**
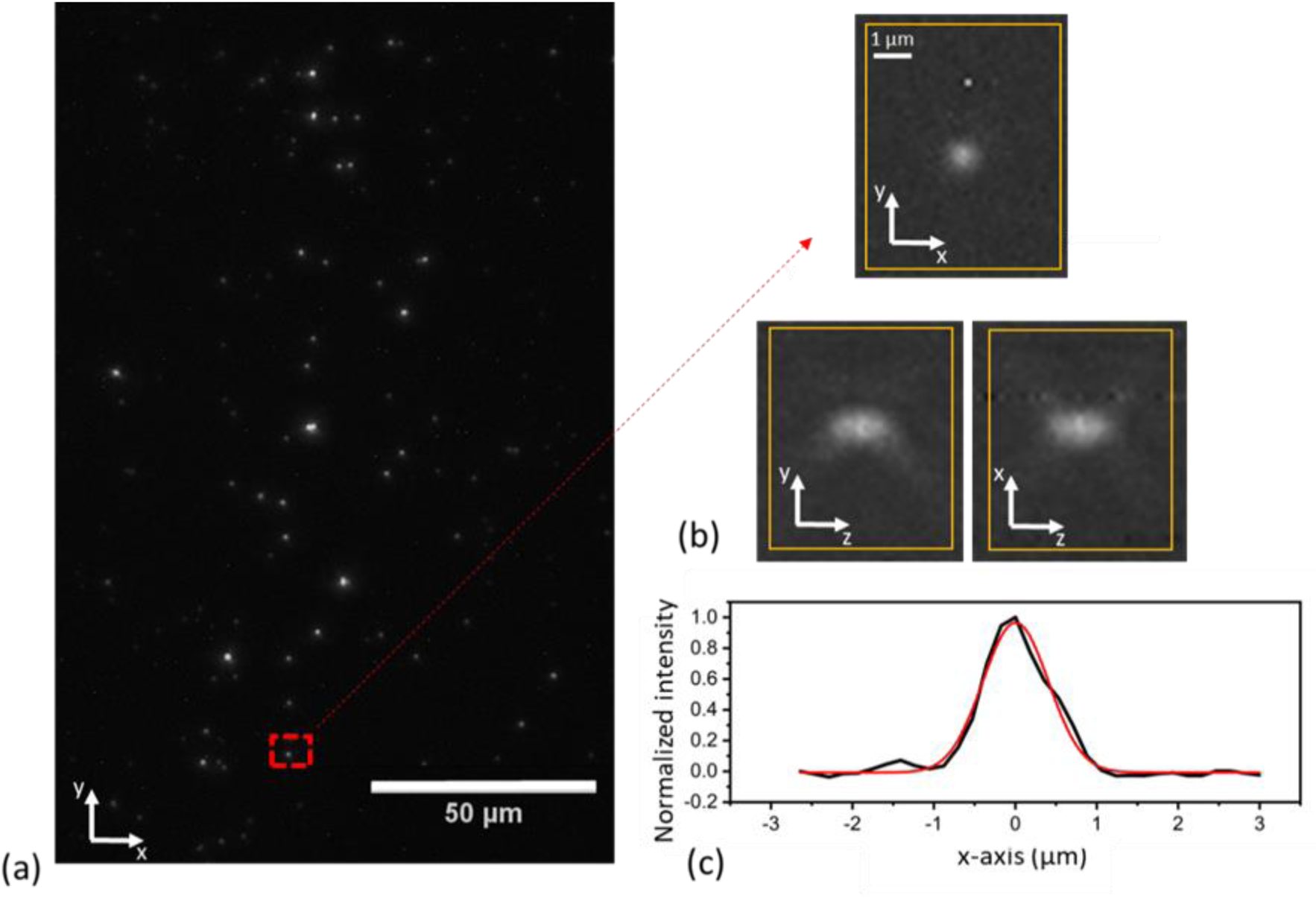
Point source imaging to evaluate the PSFs of the system. a) An exemplar maximum intensity projection of 500nm fluorescence beads at 50μm imaging depth, b) xy, xz and yz images of an exemplar bead C) cross-section of the lateral imaging PSF.

To evaluate the fluorescence point spread function of the microscope, 500 nm diameter sub-resolution nanobeads were embedded in index matching medium and fixed on a cover slip. The nanobeads were imaged close to the cover slip as well as at a depth of 57 μm (see Figure 6 (a)). Maximum intensity projections along the xy and xz and yz planes (Figure 6 (b)) show an in-plane resolution of 0.9 µm (Figure 6 (c)) and axial resolution of 13 µm in the centre of the FOV (based on 5 nanobeads measured at each location and actuation of the tunable lens to recover the PSFs for the imaging arm only), which does not degrade significantly in depth in the non-scattering medium. Images were acquired with a laser power of 0.3 mW, camera exposure time of 50 ms, axial steps of 1.8 μm but without camera binning to allow more accurate resolution estimation of the system.

### Imaging of cell slides

Imaging of fixed cells is performed with the use of fluorescent labelled bovine pulmonary artery endothelial (BPAE) cells, with F-actin stained using Alexa Fluor 488 phalloidin. Following the angled slide placement and the light-sheet generation of the bead measurements, the whole sample is scanned with 100 axial steps equal to 0.6 μm width ensuring Nyquist sampling in the z-scan. The camera exposure time is 30 ms and a 2 by 2 binning approach is taken to increase light collection as well as balance the lateral resolution close to the Nyquist sampling limit. The laser power is set at 0.3 mW, measured in the sample space. A respective tunable lens total voltage change of 0.1 V is addressed in 100 steps to match the light-sheet positions creating an imaging volume of 340 × 190 × 60 µm^3^. The optical 3D scanning is completed in a total of 4 sec per volume.

Figure 7 shows the yx (Figure 7 (a)) and zx (Figure 7 (b)) projection of the recorded 3D image, with a 10 µm thick cell slide showing a depth of over 100 µm due to the 30° angled orientation. Additionally, a plot of a selected F-actin region is shown (Figure 7 (c)) to look at the attainable resolution of the imaging this time in a more complex cell structure. In this way it is confirmed that our resolution remains below 1 μm with a clear resolving power of the F actin features of the sample.

**Figure 7.**
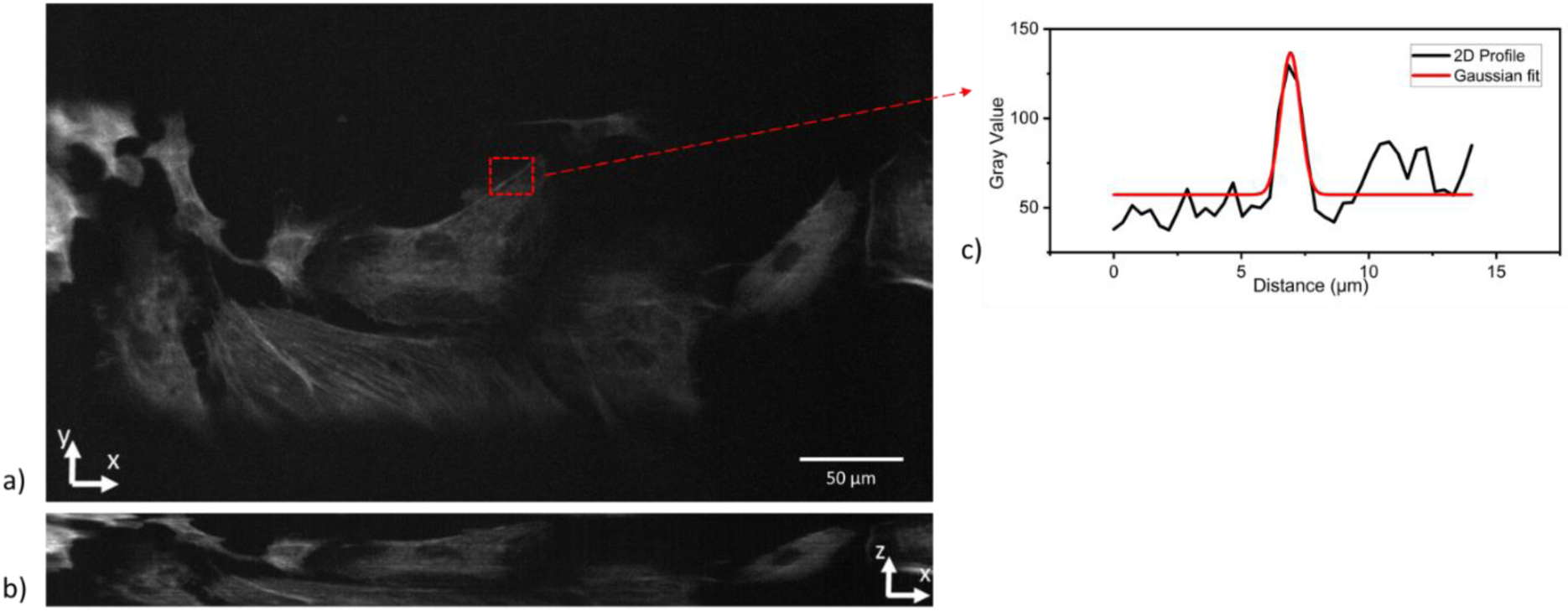
Maximum intensity projections of fixed BPAE slide with F-actin labelled with Alexa Fluor 488 (a) yz maximum projection (b) xy maximum intensity projection (c) zoom on F-actin highlighted within the intensity profiles.

## Conclusion

In this paper, we demonstrated the design and implementation of a miniaturised light-sheet microscopy system based on MEMS active optical elements. The use of micro-optical elements for beam control allows an all optical volumetric image collection, with a footprint of 20 × 28 × 13 cm^3^ allowing the potential application of the system in close controlled environments such as incubators or cell-hoods.

The presented micro-optical elements that control the optical positioning in the sample are easy to integrate into the miniaturised form factor due to their size and dual axis positioning capability. They also require low power actuation due to thin-film piezoelectric actuation, which allows simple electronics integration in the system. Additionally, they have the advantage of requiring no stage scanning which removes any motion artefacts and also allow implementation of random-access imaging. The movement settling time in the single millisecond range allows fast addressing of the imaging volume. While the presented micromirrors only show static optical scan angles of <2°, design variations will allow an increase to 10° or more, which will increase the depth of the imaging volume as only a quarter of the available tunable lens range is currently used. Implementation of further micro-optical elements can additionally allow the reduction of the axial resolution through depth scanned excitation beams synchronised with the camera acquisition.

The miniaturisation additionally leads to a system cost that is over one order of magnitude reduced compared to currently available commercial implementations. We show that this still leads to sub-micrometre lateral image resolution with volume acquisition speeds of around 0.25 vps. To improve both of these parameters, the use of more expensive microscope imaging objectives or imaging cameras is possible. We followed the aim to create a small and cost-efficient system, using economically priced objectives costing ∼£100. While a wide range of higher NA long-working distance objectives can equally be used in the presented air-coupled configuration, this would increase the price by multiple thousand pounds. Equally, an increase in imaging speed is achievable by replacing the used non-cooled camera with a higher end sCMOS camera (e.g. Photometrics Prime BSI range, Hamamatsu Orca Flash range, PCO Edge range or Andor Zyla range) to increase low light fluorescence image collection and therefore allow reduction of the camera integration time which is limiting the volume acquisition. Overall the presented system shows a good compromise of form factor, image quality and speed at an affordable price tag which will enable implementation in cost constrained settings, as well as restricted environments.

## Methods

### Custom microscope

A miniaturized LSM system as detailed in Figure 1 is designed and built. A 50 mW 473 nm solid-state laser (CNI OEM-I-473) is fibre coupled through a single-mode fibre (Thorlabs P1-460B-FC-1) and collimated with 400 μm beam diameter towards a 2-axis MEMS mirror using a focal adjustable fiber collimator package (Thorlabs CFC2-Al) and half inch silver mirror (Thorlabs ME05-P01). The 2-axis scanning MEMS mirror with 1.1 mm diameter is mounted on a small 3-axis translation stage (Thorlabs DT12XYZ/M) and mirror mount (Newport M05) to allow reflection of the excitation laser from the centre of the MEMS and accurate positioning of the vertical light sheet generation and horizontal light sheet positioning. The light sheet is generated using a sinusoidal voltage from a function generator (Rigol DG1022Z), with orthogonal positioning control achieved through a 12bit DAC and amplifier (MCP4922 and MC33072) controlled by an Arudino Due. The reflected beam enters a 4f magnifying telescope consisting of two small achromatic lenses L1 (f = 7.5 mm, Thorlabs AC050-008-A) and L2 (f = 30 mm, Thorlabs AC127-030-A). The illumination path is completed with a final achromatic focusing lens L3 (f = 7.5 mm, Thorlabs AC050-008-A) which is underfilled and has an effective NA of 0.15 and working distance of 5.2 mm. All three achromatic lenses are housed in a half inch lens tube (Thorlabs SM05L30C) with custom 3D-printed adapters to ensure accurate positioning and co-linearity between the optical elements. The orthogonal imaging beam path is designed with a 20x economy objective with 0.4 NA (Newport MVC-20X) placed at a working distance of 8.72 mm from the central illumination path focusing point. An electrical tunable lens (Optotune EL-3-10) with 3 mm aperture is positioned at the back of the objective to allow all optical synchronization of the imaging plane with the light-sheet position. The sample plane is imaged onto a 1920×1080 pixel sCMOS camera (Thorlabs CS2100M-USB) with an attached fluorescence emission filter (Thorlabs FELH0500). The imaging provides sampling of 0.36 µm/pixel with 2×2 binning, which satisfies the Nyquist criterion and is slightly oversampled. All optical components of the imaging path are housed in a lens tube (Thorlabs SM1 series) to ensure parallel optical alignment and reduce the amount of ambient light affecting the imaging outcome. Labview and the mentioned Arduino Due is used to synchronise and control the MEMS and tunable lens position as well as the image acquisition. The recorded data is analysed and processed using Fiji.

### MEMS mirror and tunable lens

The piezoelectric 2D scanning MEMS is fabricated by Stanley Electric Co., Ltd., Japan, using a 30 μm thick silicon device layer isolated by a 2 µm thick oxide from a 500 µm thick silicon handle wafer. Static tilt actuation is controlled by segmented unimorph thin-film PZT actuators (250 nm Pt/Ti bottom electrode, 4 µm PZT film, 200 nm Pt top electrode) deposited on meandering spring suspensions connecting the mirror frame to the handle wafer. An applied DC voltage will create in-plane stresses in the actuator, which translate to an out-of-plane tilt through the meandering suspension. Fast resonant tip actuation is controlled on a mirror frame with similar thin-film PZT actuators, which create a resonant scan movement around straight suspension beams connecting the mirror with the frame. The mirror has a diameter of 1.1 mm and a 100 µm thick backside reinforcement rib-structure which leads to a surface flatness of < λ /20. The static tilt actuation is controlled using an Arduino Due and a custom voltage amplifier with x 20 gain to handle the dual-side movement of the static axis through the two PZT segments. The Arduino software is integrated in a LabView GUI where the input parameter of the static axis movement can be adjusted for optimal scanning. It is found that optimal axial displacement of the axis in 100 intervals equal to 0.6 μm can be achieved with input voltage steps equal to 0.4V.

The 3 mm aperture electrical tunable lens (Optotune EL-3-10) allows positive and negative focal change by up to +-13 dioptre using a voice-coil based actuation with a control voltage of ±1 V. The lens is controlled using the Arduino Due integrated 12 bit DAC combined with a current amplifier (Texas Instruments OPA552PA) to provide up to 120 mA drive current for the lens membrane displacement. Characterization of the lens enabled focal tuning range in the microscope setup is achieved using the imaging path of the microscope setup together with a USAF 1951 target mounted on a motorized linear stage (Throlabs MTS25-Z8). Synchronization of the PZ scanning MEMS and electrical tunable lens is achieved within LabView where driving of the device during a single scan is performed in steps of 0.7 mV.

### 3D-printed prism

In order to reduce aberrations due to the angled excitation and imaging of slide-mounted samples we use a 30° prism. The prim is 3D-printed using a Formlabs Form 3 3D-printer with Formlabs Clear resin V4 as material. The prism is printed in an upright configuration with x y and z resolution equal to 25 μm. Post processing of the prim starts with a 15-minute IPA washing of the print within an ultrasound bath to remove excess uncured resin. We choose to surface-finish the prism sides using microscope slides by applying a thin layer of resin of 1 ml volume between the two and applying pressure to create a uniform coating of the prism. For this process, a standard microscope slide is cleaned with IPA and dried. Once the resin is deposited on the centre of the cleaned area, the slide is placed into a spin coater at 850 rpm for 30 seconds to ensure an evenly distributed layer of resin. The prism is pressed onto the coated surface and the slide with the prism is placed into a vacuum chamber for 20 minutes to remove potential air bubbles formed in the resin. To cure the prism side the combined elements are placed into a UV curing lamp enclosure for 60 minutes under 405nm UV LED illumination. Detaching of the prism from the microscope slide is done with the use of an accurate sharp cutting tool. The process is repeated for all three subsequent sides.

### Sample preparation

For evaluation of the system PSF, green fluorescence labelled 500 nm diameter nanobeads (PS-Speck™ Microscope Point Source Kit P7220 by Thermofisher) were used. To achieve a bead density sufficient for PSF evaluation, 0.5 μl of the stock solution were diluted by 1:10000 in ProLong™ Glass Antifade (P36984, Fisher). The antifade mounting solution was chosen to match the refractive index of the sample holder and 3D-printed prism of 1.52, in an effort to reduce aberrations between the different media. The solution is distributed into custom 3D printed mini wells, sealed over with a coverslip and cured at room temperature for 18 hours. The final test slide can be imaged in the same configuration as the fixed microscope slide samples.

All other samples are of commercial origin and bought in their imaged form.

### Image acquisition and processing

The microscope is controlled using a custom software written in LabView. Image acquisition is integrated in the synchronization algorithm and implemented in the same LabView code. The images are saved as 16-bit text files and post-processed using Fiji. Evaluation of USAF 1951 group 7 target images and 15 µm microbeads are used for calibration of the pixel/μm ratio, allowing evaluation of the FOV and imaging resolution throughout the addressable imaging space. For fluorescence imaging of the nanobeads, binning by 2×2 pixels is chosen to increase contrast and brightness, and simultaneously still maintain Nyquist sampling. Furthermore, manual contrast adjustment takes place in Fiji in order to avoid clipping and enhance image quality simultaneously for each volume.

Similar binning and processing is used for cell images where the fluorescence signal is comparably weaker. The images are grouped as 3D stacks and manual adjustment of the brightness/ contrast ratio is performed over the volume. Camera hot pixels are filtered through removing outliers with a threshold of 30 for 2 pixel radius objects. The cell scan image is presented as maximum intensity z projections for all directions.

## Data availability

All research data and materials supporting this publication can be accessed at https://doi.org/10.15129/96b96f33-0956-4621-a203-282fa799e7ed

## Acknowledgements

We acknowledge funding from the UK Engineering and Physical Sciences Research Council (grant EP/S032606/1) and UK Royal Academy of Engineering (Engineering for Development Fellowship scheme RF1516/15/8).

## Author contributions

R.B. conceived the idea, which was further developed in collaboration with S.B. and D.U. S.B. and R.B. designed and built the system. S.B. conducted the experiments and analysed the results. H.T. designed and supplied the MEMS mirror. The manuscript was prepared jointly by all authors.

